# Dynamic of Mayaro virus transmission between *Aedes aegypti* and *Culex quinquefasciatus* mosquitoes and a mice model

**DOI:** 10.1101/2022.11.20.517299

**Authors:** Larissa Krokovsky, Carlos Ralph Batista Lins, Duschinka Ribeiro Duarte Guedes, Gabriel da Luz Wallau, Constância Flávia Junqueira Ayres, Marcelo Henrique Santos Paiva

## Abstract

Mayaro virus (MAYV) is transmitted by *Haemagogus spp*. mosquitoes and has been circulating in Amazon areas in the North and Central West regions of Brazil since the 1980s, with an increase in human case notifications in the last 10 years. MAYV introduction in urban areas is a public health concern once the infection can cause severe symptoms similar to other Alphaviruses. Regarding to urban transmission, studies with *Aedes aegypti* demonstrate the potential vector competence of the species and the detection of MAYV in urban populations of mosquitoes. Considering the two most abundant urban mosquito species in Brazil, we investigated the dynamics of MAYV transmission by *Ae. aegypti* and *Culex quinquefasciatus* in a mice model. Mosquito colonies were artificially fed with blood containing MAYV and infection (IR) and dissemination rates (DR) were evaluated. On the 7^th^ post-infection day (dpi), IFNAR BL/6 mice were made available as a blood source to both mosquito’s species. After the appearance of clinical signs of infection, a second blood feeding was performed with a new group of non-infected mosquitoes. RT-q PCR and plaque assay were carried out with animal and mosquito’s tissues. We found for *Ae. aegypti* a IR of 97,5-100% and a DR of 100% in both 7^th^ and 14^th^ dpi. Regarding *Cx. quinquefasciatus*, the IR found was 13.1-14.81% and DR ranged from 60% to 80%. To evaluate the mosquito-mice transmission rate, 18 mice were evaluated (Test=12 and Control=6) for *Ae. aegypti* and 12 animals (Test=8 and Control=4) for *Cx. quinquefasciatus*. All mice bitten by infected *Ae. aegypti* showed clinical signs of infection while all mice exposed to infected *Cx. quinquefasciatus* mosquitoes remained healthy. Viremia found in those animals ranged from 2.5 × 10^8^ to 5 × 10^9^ PFU/ml. *Ae. aegypti* from the second blood feeding showed a 50% infection rate. Our study showed the applicability of an efficient model to complete arbovirus transmission cycle studies and suggests that the *Ae. aegypti* population evaluated is a competent vector for MAYV highlighting the risk of establishment of MAYV urban cycle. The mice model employed here can be used more extensively for arthropod-vector transmission studies, with laboratory and field mosquito populations, as well as with other arboviruses.

**Author summary:** Mayaro virus (MAYV) is an arbovirus maintained mostly in a sylvan cycle in South America, circulating between Haemagogus mosquitoes and wild animals. In Brazil, MAYV has been circulating in the northern region since early 80s, but a substantial increase in human cases has been reported in the past decade. MAYV infections may go undetected, as clinical symptoms are mistaken with other arboviruses already circulating in Brazil, such as dengue (DENV), Zika (ZIKV) and chikungunya (CHIKV) viruses. The introduction of MAYV in other parts of Brazil may result in a public health concern, since the virus will find all favorable conditions in urban settings: high mosquito densities, poor sanitation and uncontrolled urbanization. Therefore, we conducted a study to test the vector competence of MAYV in the two most abundant mosquito species in Brazil: *Aedes aegypti* and *Culex quinquefasciatus*. We used an animal model to analyze the dynamics between artificially-infected mosquitos and mice. We fed mosquito colonies with blood containing MAYV and on the 7^th^ day post-infection (dpi), mice were made available as a blood source to both mosquito’s species. When these mice display signs of infection, a second blood feeding was performed with a new group of non-infected mosquitoes. We found that *Ae. aegypti* mosquitoes are very competent in transmitting MAYV, while *Cx. quinquefasciatus* presented lower rates of infection and dissemination of the virus. All mice bitten by infected *Ae. aegypti* showed clinical signs of infection. On the other hand, all mice exposed to infected *Cx. quinquefasciatus* mosquitoes remained healthy. We also found a higher viremia in animals bitten by infected-*Ae. aegypti*. Overall, our study showed the applicability of an efficient model to complete arbovirus transmission cycle studies and suggests that the *Ae. aegypti* population evaluated is a competent vector for MAYV highlighting the risk of establishment of MAYV urban cycle.

## Introduction

Mayaro virus (MAYV) is an arbovirus from the *Togaviridae* family and *Alphavirus* genus that was first identified in Trinidad in 1954 (1). Since then, MAYV has been found in several countries in Latin American, mainly in regions of the Amazon area (2–5). In Brazil, the virus has been circulating for decades and recently caused outbreaks in the Northern and Central West regions of the country (6). Symptoms caused by MAYV are not specific and can range from fever, chills, gastrointestinal symptoms, retro orbital pain, myalgia and arthralgia (7,8). Similar to Chikungunya virus, MAYV causes arthralgia in 50 to 90% of patients and myalgia affects 75% of infected humans (9). In particular, arthralgia can last from months to years and severe complications such as myocarditis, hemorrhagic and neurological manifestations can also affect patients (10). MAYV surveillance in Brazil was recently intensified, mainly due to the need for differential diagnosis for Alphaviruses presenting similar clinical symptoms and have limited diagnostic laboratory facilities in many of the endemic regions (11).

MAYV has a sylvatic transmission cycle maintained between *Haemagoggus* spp. and non-human primates as vertebrate hosts. However, other species, such as from genus *Mansonia, Culex, Sabethes, Aedes* and also other vertebrate hosts (rodents and reptiles) may be involved in the transmission cycle (12–15). Once this virus has a large host range and is able to infect various vertebrates and be transmitted by different mosquito vectors, an important concern is its introduction in urban areas. Vector competence investigations have already demonstrated the ability of *Aedes aegypti* and *Aedes albopictus* to transmit MAYV (16–18). While, *Culex. quinquefasciatus* species is an inefficient vector for this virus (17). Studies with field-caught mosquitoes showed eight genus of mosquitoes naturally infected with MAYV including mosquitoes from species *Ae. aegypti, Ae. albopictus* and *Cx. quinquefasciatus* (12,19,20). Vertical transmission in *Ae. aegypti* has already been evidenced with the presence of MAYV in eggs collected in the field (21). The transmission cycle dynamics of MAYV involving different species of mosquitoes and vertebrate hosts is stills unclear. Therefore, vector competence studies addressing the whole mosquito vector-vertebrate host cycle are needed to broadly assess different mosquito vector populations and viral strains, since a set of environmental and genetic factors of viruses and vectors can alter the potential viral spread. Currently, there is no published study using vertebrate animal models focused on studying the full arthropod-vertebrate MAYV infection/transmission cycle (10).

In pathogenesis and vaccine studies, experimental animal models used for MAYV are mainly murine models in three categories: neonatal model of lethal challenge, immunosuppressed models of lethal disease and arthritis model (22–24). The increased circulation of MAYV makes it necessary to establish a new study model and with experimental focus in transmission cycle and pathogenicity related to the chronic symptoms of the infection (10). Artificially mosquitoes feeding with MAYV or mice infection is a good alternative to evaluate parameters such as vector competence and pathogenicity in mammals. However, experimental methods do not necessarily recapitulate the complexity of a blood meal in a live host, which potentially contains host immunological factors and viral antigens that can influence mosquito infection and virus transmission (25–28). In vector competence studies are also gaps and lack of standardization in relation to the methods and nomenclatures used in the assays (29). Investigation with an animal model and stages of transmission cycle should be performed with the aim of improving our understanding of pathogen-vector-host interaction dynamics.

Considering the two most abundant species in the urban environment in Brazil, the aim of the present study was to evaluate the transmission dynamics of MAYV by *Ae. aegypti* and *Cx. quinquefasciatus* from Recife-PE using an animal model (mosquito-host-mosquito). The focus of the study was to characterize the virus-vector-host interaction for MAYV, once a complete transmission model has not yet been described for MAYV and the imminent risk of emergence of the virus through species such as *Ae. aegypti*.

## Material and methods

### Cells and viral strain

VERO CCL81 cells were grown in Minimum Essential medium (MEM; Gibco, catalog #61100087, USA) supplemented with 10% fetal bovine serum (FBS; Gibco, catalog #A31608, USA), and 1% penicillin/streptomycin (Gibco, catalog #15140-122, USA) at a 37°C + 5% CO_2_ incubator. MAYV strain used in mosquito’s infection was MAYV/BR/Sinop/H307/2015 (MH513597) derived from human isolate kindly provided by Dr. Roberta Bronzoni from the Federal University of Mato Grosso (UFMT-Sinop). Viral stock was prepared by inoculation on to an 80-90% confluent monolayer of VERO cells. Virus culture were harvested at 24 hours after infection, centrifuged at 1,200 rpm for 10 minutes and supernatants were transferred to a new cryotube and then stored at −80°C until use. Prior to mosquito infection, virus titer was calculated via plaque assay and reached 10^8^ plaque-forming units per milliliter (PFU/mL).

### Mosquitoes and animals

Mosquitoes’ colonies used in the present study were derived from two natural populations collected in Recife/PE, Brazil: RecLab (*Ae. aegypti*) and CqSLab (*Cx. quinquefasciatus*). These mosquito colonies have been maintained at standard conditions of 26±2°C, 65%–85% relative humidity, 10/14 light/dark cycle in the Entomology department at Aggeu Magalhães Institute (AMI) for several generations (Melo-Santos et al. 2010; Amorim et al. 2013).

Concerning animal models for virus transmission, we used IFNAR Bl/6 (-Ifnar1^tm1.2Ees^/J) mice strain which was provided by the Jackson Laboratory Repository (JAX stock #028288, USA) to the Central Animal Facility of the University of São Paulo, School of Medicine, in Ribeirão Preto/SP, Brazil. This center kindly provided the animals to the AMI Animal Facility installation with the necessary sanitary and genetic certification.

### Mosquitoes artificial feeding and vector competence evaluation

Seven-to-ten-days-old mosquitoes from RecLab and CqSLab colonies were separated in different plastic cages named test and control groups for each species. Test group contained 200 females and 50 males, while control group contained 100 females and 30 males. Twenty-four hours prior to artificial blood feeding, both species were sugar starved. Artificial blood feeding assay was carried out in a NB2 (Biosafety Level 2) and after that all the plastics cages were maintained in containment cages. Experiments were conducted using a fresh MAYV grown in VERO cells at multiplicity of infection of 0.1 collected after 24 hours of inoculation for the test group and uninfected cultured cells for the control group. The cell culture flasks were frozen at −80°C, thawed at 37 °C and then mixed with defibrinated rabbit blood in a 1:1 proportion and ATP (Sigma-Aldrich, catalog #A2383, USA) in a 3 mM final concentration. Briefly, blood mixture was offered in 4 cm diameter petri dishes containing a triple membrane of PARAFILM® M (Sigma-Aldrich, catalog #P7543, USA) on top of the cages at 37 ºC by using heat packs for one hour. Fully engorged females were cold anesthetized, transferred to a new cage and maintained in the infection room for 14 days’ post infection (dpi). Blood mixtures used in the assays as well as fully engorged females were collected and processed as artificial feeding check samples.

In order to investigate vector competence of *Ae. aegypti* and *Cx. quinquefasciatus* previously fed with MAYV. Fifteen females for each species were collected at the 3^rd^, 7^th^ and 14^th^ days post infection (dpi). Vector competence status was estimated by the detection of MAYV in dissected midguts (MID) and salivary glands (SG) at different time points (Fig 1A). Tissues were collected in a petri dish containing cold PBS 1X (Thermo Fisher Scientific, catalog #J75889, USA) with forceps and visualization in a stereomicroscope (LABOMED, USA). Dissection was performed quickly in ice and before every mosquito’s dissection, all instruments were cleaned with 70% ethanol. Immediately after dissection, tissues were individually transferred to a 1.5 ml DNase / RNAse-free microtube containing 300 μl of mosquito diluent (32) and stored at -80°C until further use.

**Figure 1:**
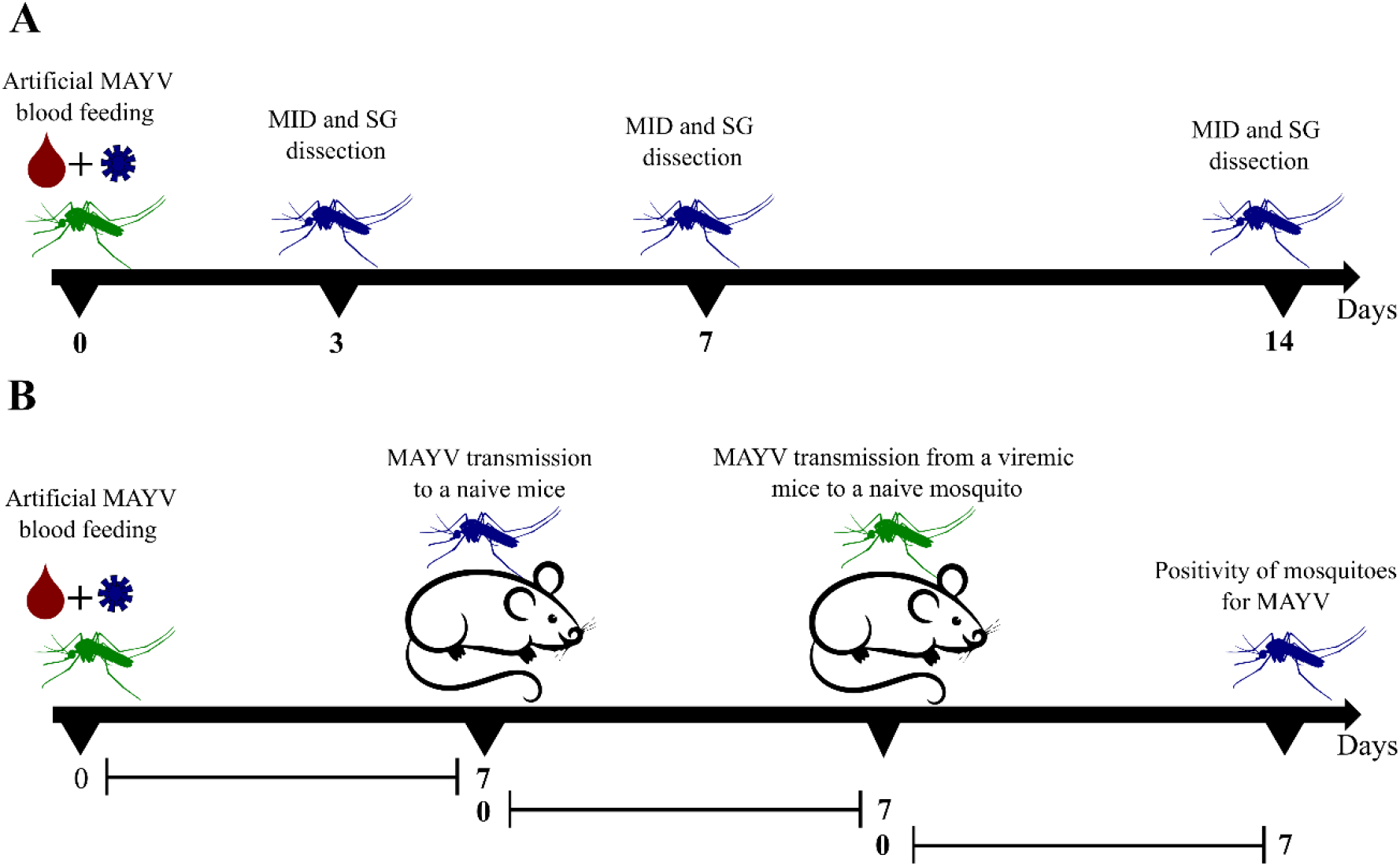
Illustration of vector competence study design and Mayaro virus transmission cycle model in mice. **(A)** Vector competence study design in a time line. **(B)** Mayaro transmission cycle model in a time line. **MID-** Midguts; **SG-** Salivary glands and **MAYV-**Mayaro virus. Illustration was created by the authors using Inkscape software v 1.2.

### MAYV transmission cycle model in mice

In order to determine whether viral load detected in mosquitoes is sufficient to infect naïve mice and then to new susceptible mosquitoes (complete transmission cycle), transmission dynamics assays were performed with IFNAR/Bl6 mice aged between 2-4 weeks, males and females, separated in micro-isolators into test groups comprised of four animals and control groups comprised of two animals. Animals were kept at 22±2 °C, 50%–60% relative humidity, 12/12 light/dark cycle and were acclimated for at least 72 hours before use. The time point (3, 7 or 14 dpi) for the analysis of animal transmission by the mosquitoes infected with MAYV was evaluated after vector competence assays. At the chosen time point when mosquito colonies showed high dissemination rates, both animal groups (test and control) were weighed, anesthetized with the association of Ketamine (100mg/kg) (Syntec, Brazil) and Xylazine (10mg /kg) (Syntec, Brazil) and then placed on the surface of plastic cages containing mosquitoes (10-15 females per mice) for 20 minutes. During the procedure, animals were monitored for any signs of suffering. After the blood meal, animals were monitored daily for seven days to assess weight loss, clinical signs and mortality rate (Figure 1B). Clinical signs had numerical values assigned according to the following score: Isolation-1, ruffled hair-1, mild lethargy-1, curved posture-2, moderate lethargy-2, weight loss (≥20%)-2, foot edema-2, severe lethargy-3 and diarrhea-3.

At the moment that signs of infection were identified (viremia period considered with score ≥3), animals from the test group were again anesthetized to be available as a blood source for a new group of naive mosquitoes (10-15 females per mice) according to the protocol described above. After the blood meal in infected mice, mosquitoes were kept in the infection room for another 7 days and then collected, anesthetized at -20 ºC, and individually stored in 1.5 ml RNAse/DNAse free tubes containing 300 μl of diluent (32) (Fig 1B).

With regards to the animals after the second blood meal with naïve mosquitoes, it was collected whole blood and euthanasia was carried out by cardiac puncture. After the procedure, animals were dissected to collected brain, liver and gonads from control and test groups using sterile surgical scissors and forceps. Tissues were stored in 2 ml RNAse/DNAse free tubes containing 500 μl of diluent (32). Blood was centrifuged for 10 minutes at 4,000 rpm and serum transferred to a new 1.5 ml RNAse/DNAse free tubes.

### Ethics Committee

All animal experimental procedures were approved by the Ethics Committee for Animal Use of the AMI (CEUA/IAM 166/2021) and followed the guidelines of the National Council for the Control of Animal Experimentation (CONCEA).

### RNA isolation and RT-qPCR

Mosquito and mice tissues were individually homogenized as described in Barbosa *et al*. (2016). RNA extraction was performed with TRIzol® reagent (Invitrogen, catalog#15596026, USA) as described in Guedes *et al*. (2017). RT-qPCR reaction was carried out using the QuantiNova Kit Probe RT-PCR Kit (Qiagen, catalog #208354, Germany), using primers and probe described in Naveca *et al*. (2017). Reactions were performed in a final volume of 10 μl with the following final concentrations: QuantiNova Probe RT-PCR Master Mix (1x), QuantiNova ROX (1x), QuantiNova RT Mix (1x), 800 nM of primers and 100 nM of probes. RT-qPCR reactions were performed in a QuantStudio 5 System (Applied BioSystems, USA) under the following conditions: 45 °C for 15 minutes, 95 °C for 5 minutes, followed by of 45 cycles of 95 °C for 5 seconds and 60 °C for 45 seconds. All samples were tested in duplicates, with negative reaction control (all reagents except RNA), negative control from RNA isolation and positive control (standard curve). The standard curve used was synthesized by *in vitro* transcription using MEGAscript T7 kit (Ambion, catalog #AM1334, USA) and was quantified in Nanodrop 2000. RNA concentration was converted in RNA copy number as described in Kong *et al*. (2006). Results were analyzed using QuantStudio Design and Analysis Software v.1.3.1 with automatic threshold and baseline. Samples with Cycle quantification (Cq) values ≤ 38.5 in both duplicates were considered as positive.

### Plaque assay

Viral titer in mosquito and mice tissues was calculated using plaque assay. Initially the 24-well plates containing 3 × 10^5^ cells/ml in MEM supplemented with 10% FBS were prepared and incubated at 37 °C + 5% CO_2_. After 24 hours, the medium was discarded and inoculated 50 μl of a serial dilution (10^−1^ to 10^−10^) of each sample tissue in duplicate and incubated for 30 minutes at 37 ºC + 5% CO_2_ for adsorption. After incubation, 500 μl of MEM semi-solid (3% carboxymethylcellulose) were added. After 48 hours of incubation at 37 °C + 5% CO_2_ the medium was removed by inversion and cell monolayer was fixed with 8% formalin solution (Sigma-Aldrich, catalog #A252549, USA) for one hour and 20 minutes and then revealed with crystal violet 0.04% (Sigma-Aldrich, catalog #6158, USA) for plaque visualization and counting.

### Data Analysis

The infection rate (IR = positive midguts / midguts analyzed) and dissemination rate (DR = positive salivary glands/positive midguts) were calculated for each species at different time points. Logistic regression, X^2^-test and Fisher’s Exact tests were used to test differences in both the IR and SR within the two species. Wilcoxon test was used to compare MID and SG values. One-way analysis of variance (ANOVA) followed by Tukey’s multiple comparisons test was used for mice tissues analysis. Analyzed results were considered significant when p value <0.05. All statistical tests and graphics were performed with R software (R DEVELOPMENT CORE TEAM) and Graph Pad Prism 9 software (Graph Pad, San Diego, CA, USA).

## Results

### Vector competence assays

To evaluate vector competence of *Ae. aegypti* (RecLab) and *Cx. quinquefasciatus* (CqSLab) colonies for MAYV, three independent artificial blood feeding experiments were performed for each species. The titer of the initial mixture (MAYV culture + blood) before artificial blood feeding ranged from 2.5×10^6^ to 1.5×10^7^ PFU/mL. After blood feeding, one mosquito from each group was collected to evaluate viral load decay, and the titer after 60 minutes ranged from 2,5 × 10^4^ to 1,25 × 10^5^ (Table 1). MID and SG of 135 females of *Ae. aegypti* and 100 females of *Cx. quinquefasciatus* were analyzed (Table 2). On the 3^rd^, 7^th^ and 14^th^ dpi tissues were dissected and tested by RT-qPCR to verify the presence and to quantify MAYV. For *Ae. aegypti* the IR found was 97.7% (3^rd^ dpi), 100% (7^th^ dpi), 100% (14^th^ dpi) and the DR reached 100% in both 7^th^ and 14^th^ dpi. Regarding *Cx. quinquefasciatus*, the IR found on the 3^rd^, 7^th^ and 14^th^ dpi was 14.28%, 13.1% and 14.81%, respectively. In CqSLab colony, even with a low IR, ten positive salivary glands were detected, which generated a DR that varied between 60% and 80% on the time points (Fig 2).

**Table 1:** Mayaro viral titer used in artificial blood feeding.

| Assay | MAYV Titer<br>(PFU/mL) | MAYV Titer in mosquito after blood meal (PFU/mL) |  |
| --- | --- | --- | --- |
|  |  | RecLab | CqSLab |
| 1 | $2.5 \times 10^6$ | $2.5 \times 10^4$ | $1 \times 10^5$ |
| 2 | $1.5 \times 10^7$ | $7.5 \times 10^4$ | $1 \times 10^5$ |
| 3 | $1.25 \times 10^7$ | $7.5 \times 10^4$ | $1.25 \times 10^5$ |

**Table 2:**
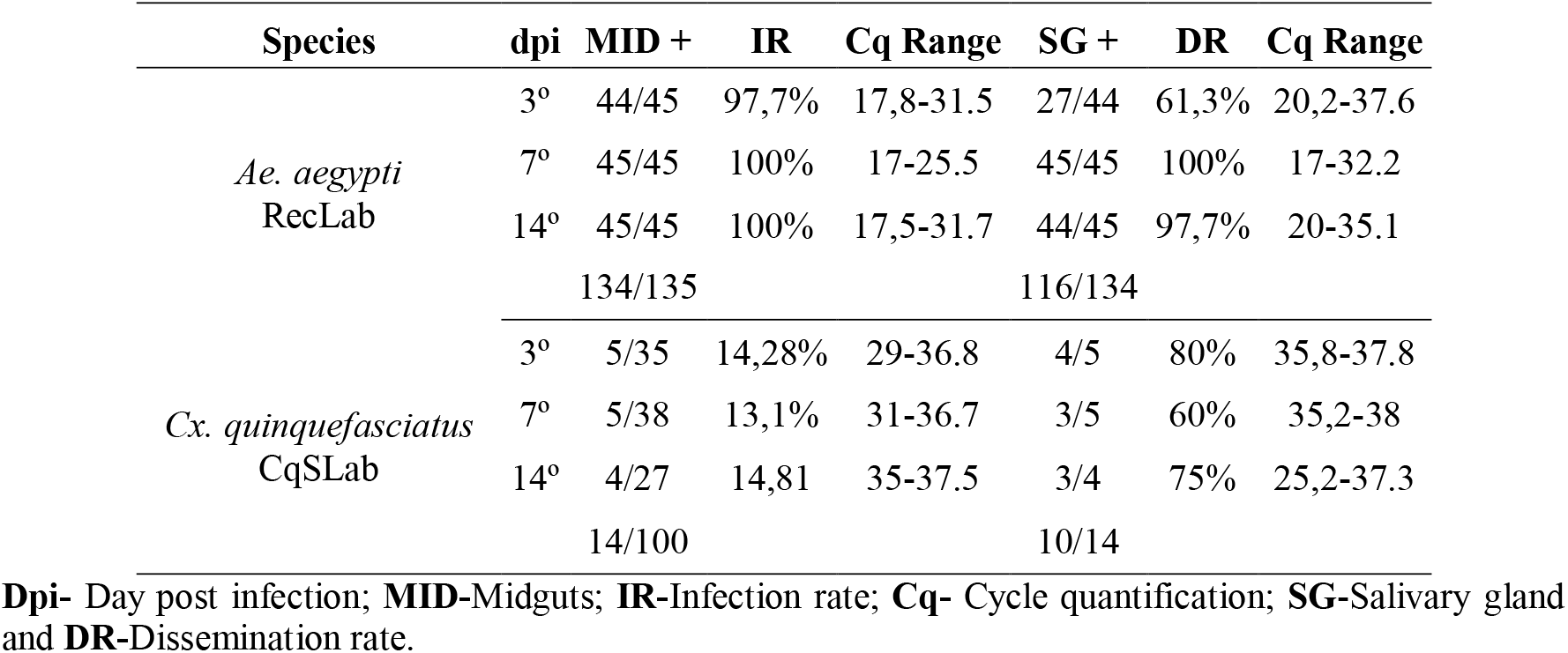
Description of positive samples, infection rate and dissemination rate for Mayaro virus from *Aedes aegypti* (RecLab) and *Culex quinquefasciatus* (CqSLab) after artificial blood feeding.

| Species | dpi | MID + | IR | Cq Range | SG + | DR | Cq Range |
| --- | --- | --- | --- | --- | --- | --- | --- |
| *Ae. aegypti*<br>RecLab | 3° | 44/45 | 97,7% | 17,8-31.5 | 27/44 | 61,3% | 20,2-37.6 |
|  | 7° | 45/45 | 100% | 17-25.5 | 45/45 | 100% | 17-32.2 |
|  | 14° | 45/45 | 100% | 17,5-31.7 | 44/45 | 97,7% | 20-35.1 |
|  |  | 134/135 |  |  | 116/134 |  |  |
| *Cx. quinquefasciatus*<br>CqSLab | 3° | 5/35 | 14,28% | 29-36.8 | 4/5 | 80% | 35,8-37.8 |
|  | 7° | 5/38 | 13,1% | 31-36.7 | 3/5 | 60% | 35,2-38 |
|  | 14° | 4/27 | 14,81 | 35-37.5 | 3/4 | 75% | 25,2-37.3 |
|  |  | 14/100 |  |  | 10/14 |  |  |

**Dpi-** Day post infection; **MID-**Midguts; **IR-**Infection rate; **Cq-** Cycle quantification; **SG-**Salivary gland and **DR-**Dissemination rate.

**Figure 2:**
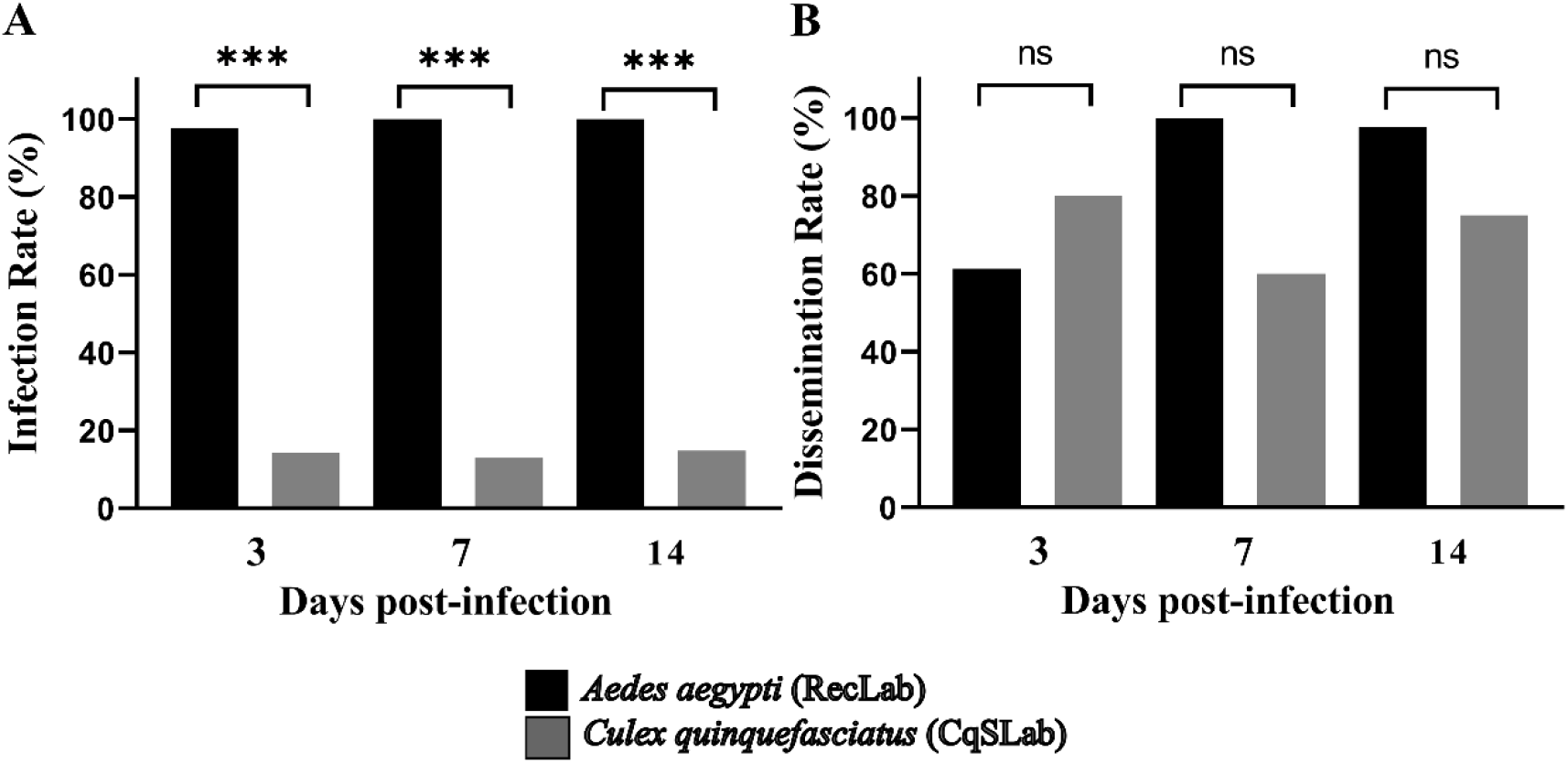
Infection and dissemination rates of *Aedes aegypti* (RecLab) and *Culex quinquefasciatus* (CqSLab) mosquitoes experimentally fed with blood containing Mayaro. **Dpi**- Day post infection; **ns**- No significant. Statistical analysis was performed using R software (R DEVELOPMENT CORE TEAM) (***P<0.001).

A total of positive MID and SG are described in table 2 and statistical analyzes demonstrated differences between values obtained for the two species and different organs. Regarding to Cq values, RecLab colony showed lower Cq values in MID (ranged from 17 to 31.7) when compared to CqSLab (ranged from 29 to 37.5) (Fig 3A). Comparing SG, RecLab showed also a lower Cq values (ranging from 17 to 37.6) when compared to CqSLab (ranging from 25.5 to 38) (Fig 3B). In terms of RNA copies number/mL, the quantification showed the same pattern on the MID samples of both species, however the SG quantification showed a difference only at the 7º dpi (Fig 3C and 3D). Plaque assays were performed with 12 GS samples from RecLab (Cq range from 17.2 to 37.5) and samples with Cq <30 showed plaque forming units. All CqSLab MID and SG samples (N=24) were submitted to plaque assay and only one SG sample (Cq=25.2) formed plaque and showed a titer of 1.25×10^4^ PFU/mL.

**Figure 3:**
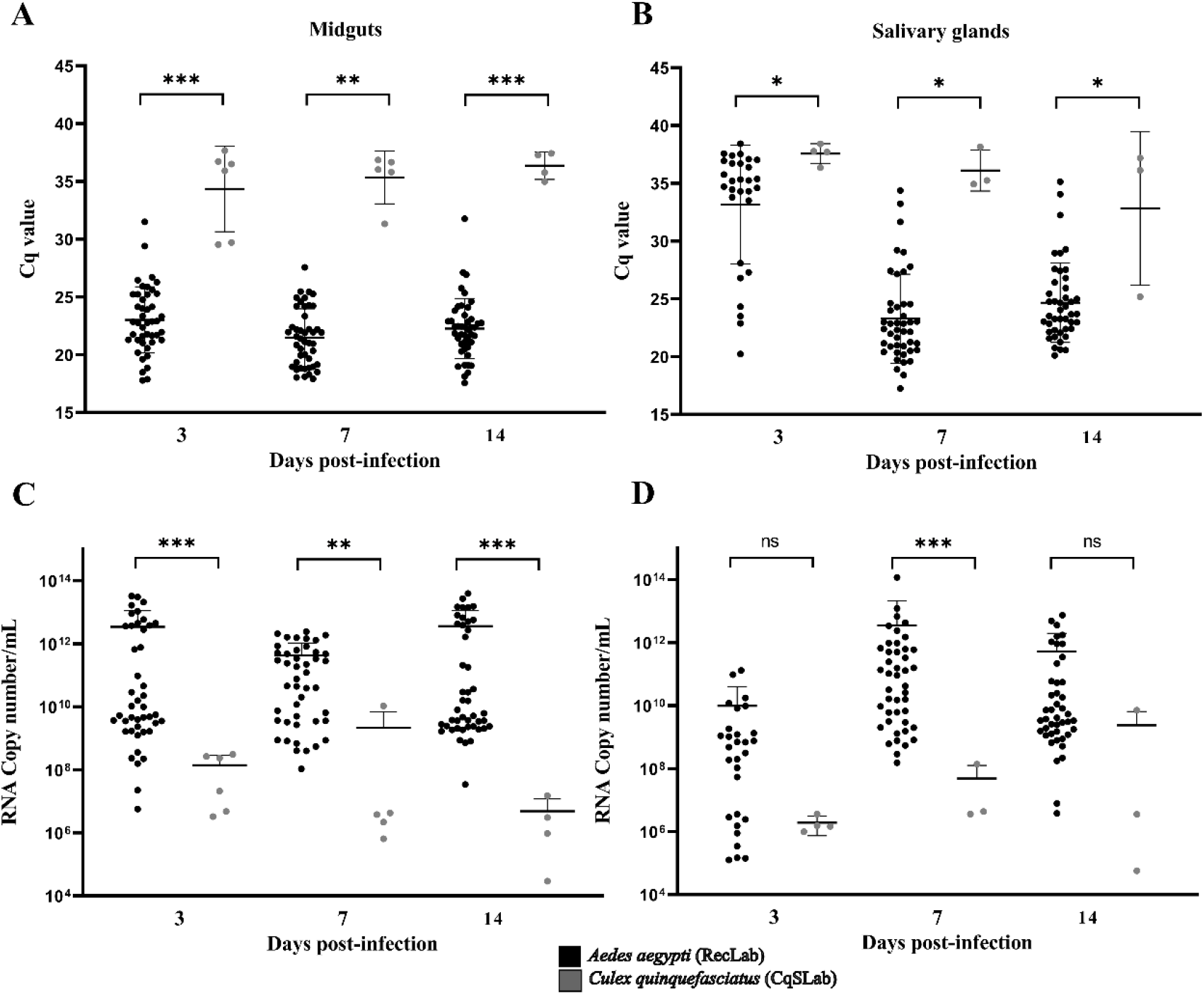
Midguts and salivary glands of *Aedes aegypti* and *Culex quinquefasciatus* positives by RT-q PCR for Mayaro. **(A-B)** Cq values in the midguts and salivary glands of positive *Aedes aegypti* and *Culex quinquefasciatus* mosquitoes experimentally fed with blood containing MAYV; **(C-D)** Quantification of RNA viral copy numbers in the positive midguts and salivary glands of *Aedes aegypti* and *Culex quinquefasciatus* mosquitoes experimentally fed with blood containing MAYV; **Dpi-** Day post infection; **ns-** No significant. Statistical analysis was performed using R software (R DEVELOPMENT CORE TEAM) (*P<0.05, **P<0.01 and **P<0.001).

### Mayaro virus Transmission cycle in mice

After analysis of infection and dissemination rates in both mosquito species, we followed up MAYV transmission rate to mice on 7^th^ dpi using RecLab colony as a reference, once this specie displayed the best results for virus replication in salivary glands. Cq values and RNA copy number/mL in *Ae. aegypti* showed higher amount of virus at 7^th^ dpi (Fig 4A an 4B).

**Figure 4:**
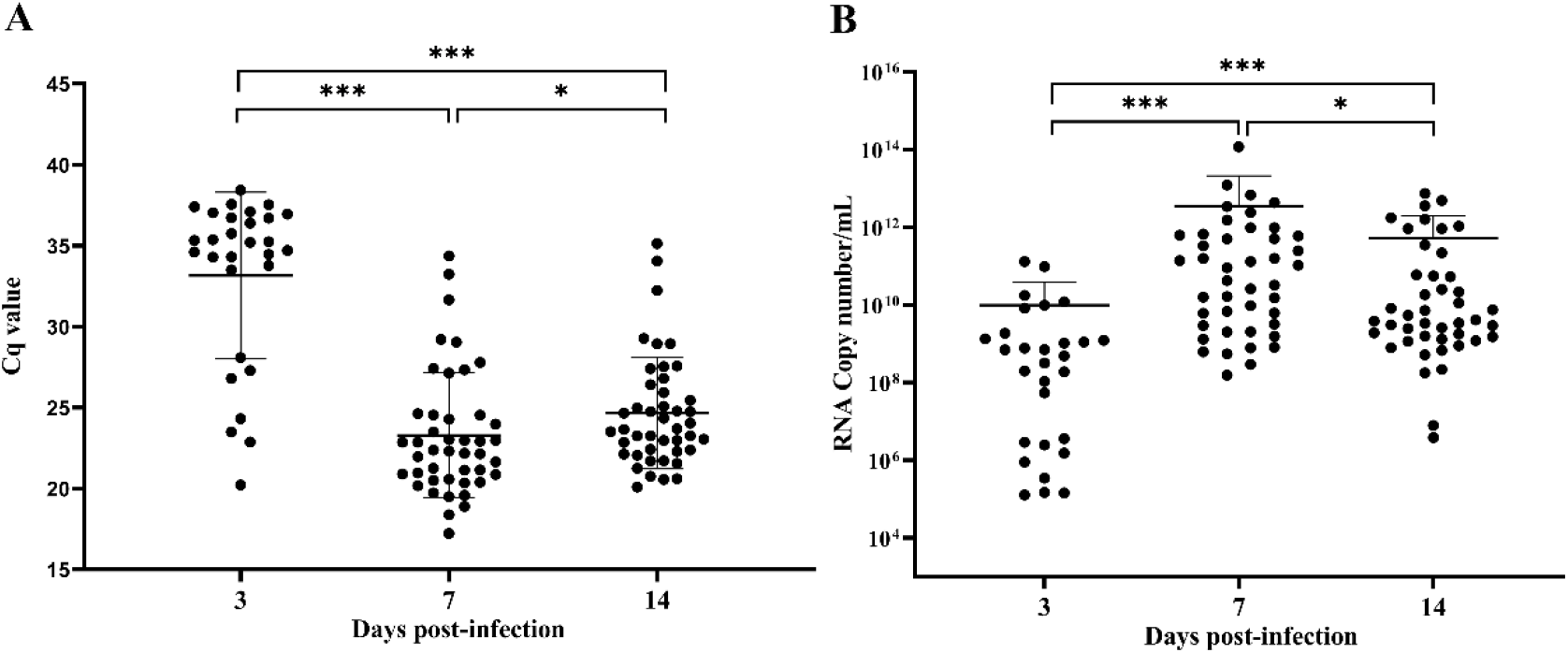
*Aedes aegypti* positive salivary glands by RT-q PCR for Mayaro virus. **(A)** Cq values of positive salivary glands samples of *Aedes aegypti* mosquitoes experimentally fed with blood containing Mayaro. **(B)** RNA copy number of positive salivary glands samples of *Aedes aegypti* mosquitoes experimentally fed with blood containing Mayaro. **Dpi-** Day post infection. Statistical analysis was performed using R software (R DEVELOPMENT CORE TEAM) (*P<0.05, **P<0.01 and **P<0.001).

Three independent experiments were performed for MAYV transmission in an animal model. A total of 30 animals was used in the experiments: 12 animals from the test group and six animals from the control group with RecLab mosquitoes and eight animals of the test group and four animals of the control group with CqSLab mosquitoes (Table 3). Eight mice infected by *Ae. aegypti* demonstrated signs of infection on the 3^rd^ dpi and, in addition four animals died on the 4^th^ dpi. Viral load was calculated in infected mice and ranged from 2.5×10^8^ to 5×10^9^ PFU/mL (Table 3). Animals presented a classic symptom of Alphaviruses, foot edema, as well as several other signs of infection (Fig 5A and 5B). The score observed in mice are described on table 3. Regarding CqSLab experiments, zero animals showed sign of infection as well as controls groups.

**Table 3:**
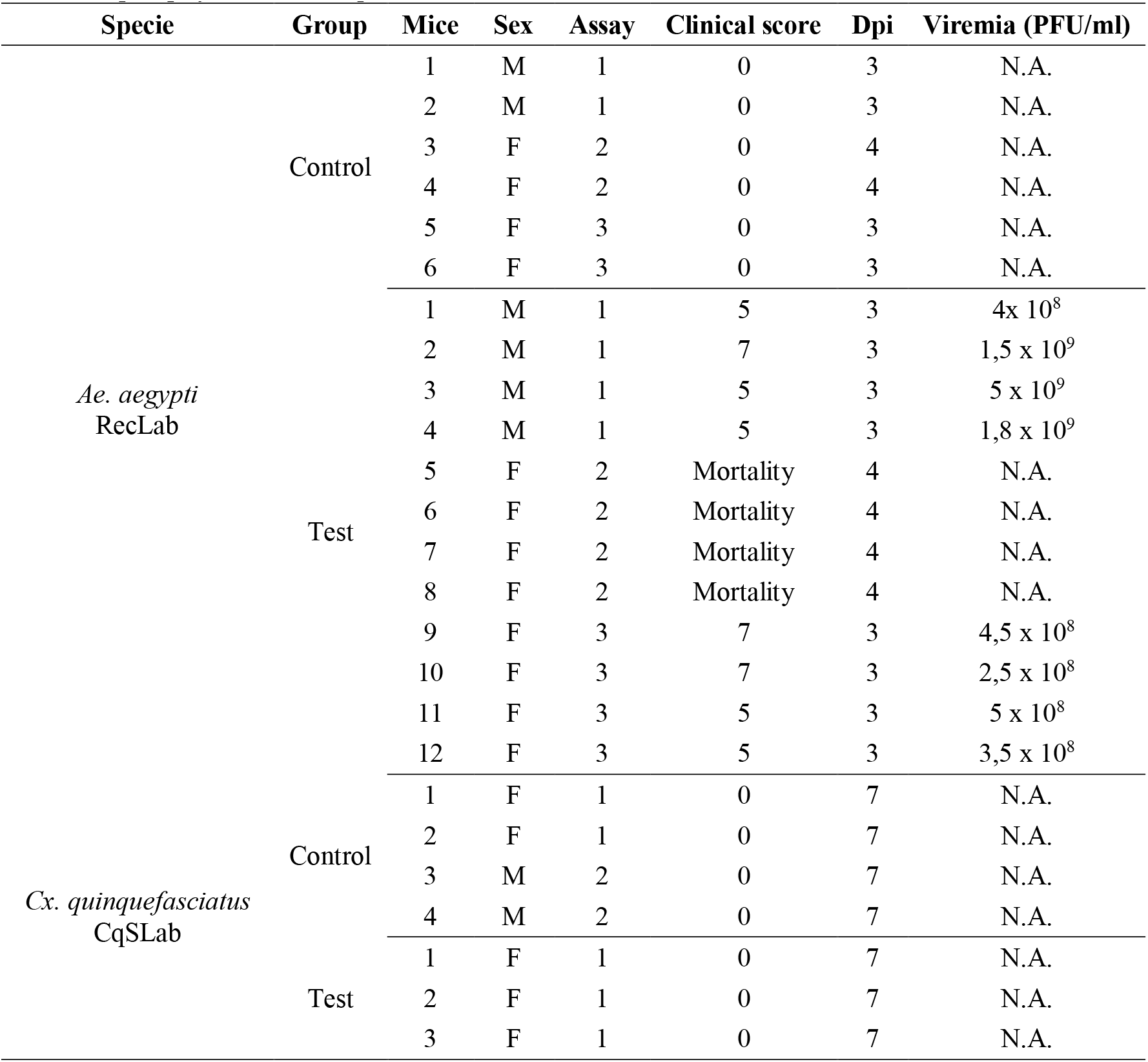
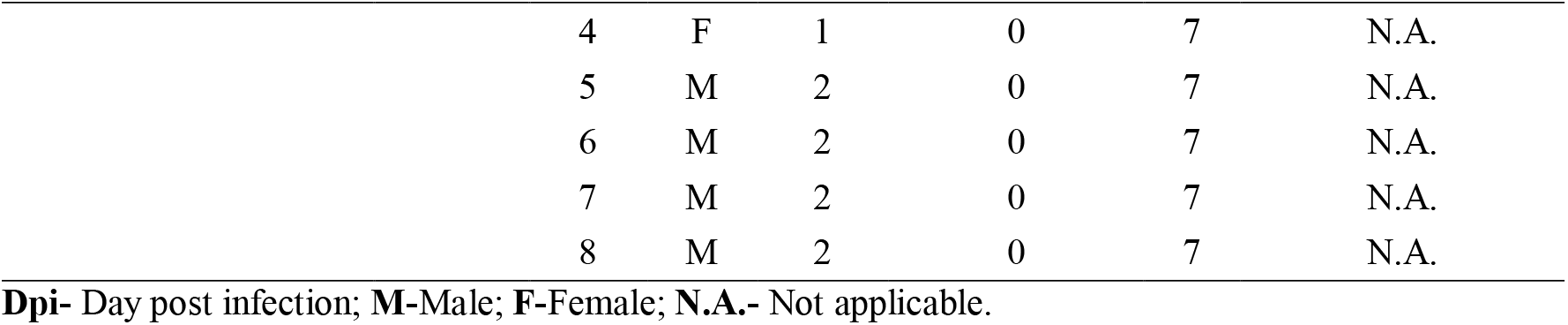
Description of IFNAR-1 mice used in Mayaro transmission cycle experiments with *Aedes aegypti* and *Culex quinquefasciatus* mosquitoes.

**Figure 5:**
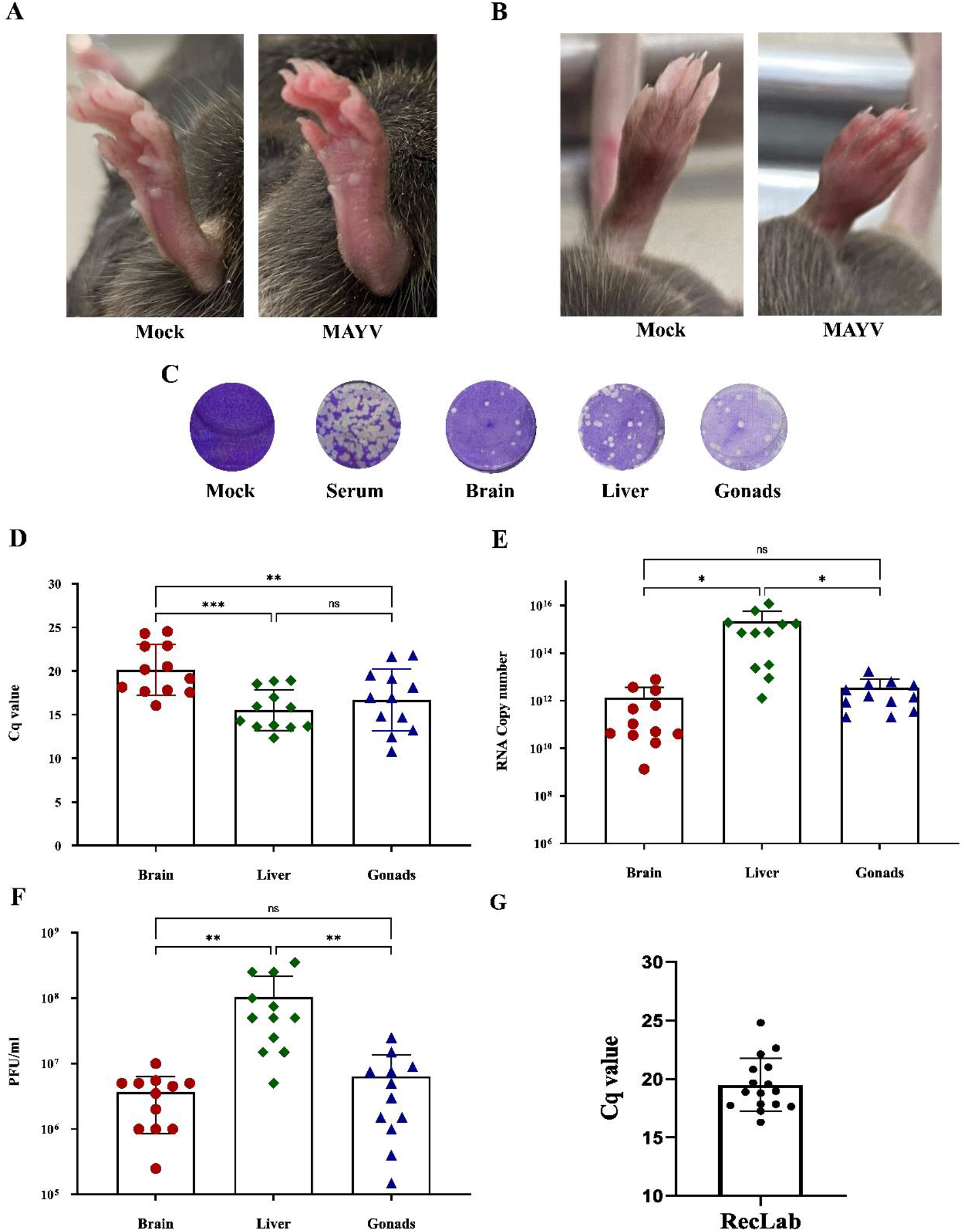
Animal experimentation panel with results of IFNAR-^1^mice infected with Mayaro and mosquitoes *Aedes aegypti*. **(A)** Foot edema presented in mice hind foot after Mayaro infection and Mock hind foot showed in bottom view; **(B)** Foot edema presented in mice hind foot after Mayaro infection and Mock hind foot showed in top view; **(C)** Visualization of plaque form unit in different mice samples at 10^−5^ dilution in VERO cell plaque assay. **(D)** Cq values of mice tissues samples infected with Mayaro after *Aedes aegypti* blood feeding. **(E)** RNA copy number of mice tissues samples infected with Mayaro after *Aedes aegypti* blood feeding. **(F)** Cq values of mice tissues samples infected with Mayaro after *Aedes aegypti* blood feeding. **(G)** Cq values of *Aedes aegypti* whole body samples after second blood feeding in mice infected with Mayaro. **ns-** No significant. Statistical analysis was performed using Graph Pad Prism 9 (*P<0.05, **P<0.01 and **P<0.001).

Fifteen females for each species Brain, liver and gonads tissues of all animals were collected, including those that succumbed to the infection and died. In summary, all organs that were processed by RT - qPCR of the control groups were negative, in CqSLab test group, out of 24 tissue samples collected, only one brain was positive with Cq value of 32.9. Animals of RecLab test group showed MAYV presence in all tissues. Organs of the infected animals were also assayed by plaque assay and all samples presented plaque forming units (Fig 5C). Cq values ranged 17.5 to 24.3 (brain), 12.3 to 18.5 (liver) and 10.7 to 21.8 (gonads). Regarding viral load (PFU/ml) titers ranged 2.5 × 10^5^ to 1 × 10^7^ on brain, 5 × 10^6^ to 3.5 × 10^8^ on liver and 1.5 × 10^5^ to 2.5 × 10^7^ on gonads. Statistical analysis was performed on Cq values, RNA copy number and viral load (PFU/ml), and with Cq values, results showed that the brain was the organ with the highest Cq value when compared to the other two organs (Fig 5D). When we looked at the quantification of RNA copy number/mL, we observed that the liver was the tissue with the highest RNA copy number /mL when compared to brain and gonads (Fig 5E). In the analysis of viral load (PFU/mL) a virus tropism for liver samples was also visualized (Fig 5F).

Concerning MAYV incubation period in mice, at 3^rd^ and 7^th^ dpi, animals of the test group were available as a blood source for RecLab and CqSLab mosquitoes, respectively. At day 7 after blood feeding on mice, all mosquitoes were collected and 30 individuals (whole body) were assayed by RT-qPCR. As a result, concerning *Cx. quinquefasciatus* no mosquito was found positive. Regarding *Ae. aegypti* mosquitoes 15 out of 30 (50%) were positive for MAYV and Cq values ranged from 17.6 to 24.8 (Fig 5G).

## Discussion

Arbovirus are maintained in nature through sylvatic and urban transmission cycles involving hematophagous invertebrate vectors (mainly mosquitoes and mites) and a range of vertebrate hosts. These viruses are responsible for outbreaks and epidemics all over the world. Mayaro virus is considered an emerging arbovirus and increasing number of human’s cases has been reported in several Latin American countries (10,12). Currently, there is no large scale and ongoing urban transmission cycle of this virus although it may be eminent in the next years. Nonetheless, there is an absence of pathogen-vector-host interaction studies for MAYV, highlighting the urgent need of such studies for viruses that already have been detected in the human population and may be one step behind of adaptations that will allows large scale sustained urban transmission in developing countries that lack resources for arbovirus transmission control. Here, we demonstrated a complete model of MAYV transmission (mosquito-host-mosquito) with two mosquito species, which are very abundant in the urban environment (*Ae. aegypti* and *Cx. quinquefasciatus*). We showed high infection and dissemination rates for *Ae. aegypti* and low IR and high DR for *Cx. quinquefasciatus*. After analysis of vector competence of both species, mosquitoes were used to test transmission of MAYV to naive mammal’s host. *Ae. aegypti* were able to transmit and infect 100% of the mice available for feeding, and clean mosquitos of this species were also able to become infected and maintain a 50% infection rate after seven days after blood feeding on MAYV infected mice.

Regarding the vector competence data, we have observed a high IR (>97%) for *Ae. aegypti* and DR (>61%), in line with findings by Kantor *et al*. (2019) using *Ae. Aegypti* populations from the USA (>80%) and Long *et al*. (2011) that used *Ae. aegypti* from Peru (>80%). Pereira *et al*. (2020) performed experimental infection assays with field populations of *Ae. aegypti, Ae. albopictus* and *Cx. quinquefasciatus* from Minas Gerais and detected infection rates for the former species of 57.5% and 61.6 for *Ae. albopictus*. These results are similar to those observed in the present study. Regarding *Cx. quinquefasciatus*,the IR found was 2.5%, much lower when compared to our estimates of 13-15%. While the DR for *Cx. quinquefasciatus* found in the cited study was zero, our data showed 10 positive salivary glands for MAYV, which generated DR of 60 to 80%. Although, when *Cx. quinquefasciatus* samples were inoculated into VERO cell monolayers, only one sample formed plaques, likely indicating low proportion of infectious particles in the sample that resulted in limited active viral replication.

The animal model proposed here was never described before for MAYV, once there are no vector competence studies that showed the complete transmission cycle for MAYV. We have demonstrated the ability of *Ae. aegypti* to transmit MAYV to a 100% of susceptible mice and then the ability of those viremics animals to transmit the virus to mosquitoes after 3 days. Some studies used saliva collection as a parameter of virus transmission (17,39). However, the detection of viruses in saliva is not an effective method, in this approach there is a methodological difficulty in confirmation of saliva collection and processing a small amount of saliva samples. In the literature we can find data of animal experimentation to evaluate virus transmission, but without vector competence evaluation and without a second blood meal as demonstrated by our investigation. Some studies addressed vector competence with different methodological approaches, Secundino *et al*. (2017) evaluated the transmission of Zika (ZIKV) by *Ae. aegypti* to mice ear, with the analysis of the presence of ZIKV in the tissue. Long *et al*. (2011) used vertebrate hosts to assess the transmission of MAYV by *Ae. aegypti*. Hibl *et al*. (2021) developed a study with a new model of infection that included the natural vectorial transmission of CHIKV by *Ae. aegypti* to mice that containing components of the human immune system and Weger-Lucarelli *et al*. (2019) also used an animal model with transmission by mosquitoes bites and focused on diet and obesity during MAYV infection.

Multiple animal models have been used to study the pathogenesis and tropism of Alphaviruses, such as CHIKV and MAYV (43,44). We demonstrated classic signs of infection like footpad edema and a short incubation period (3-4 days after infection) in the animal model used in this study, which corroborates with Santos *et al*. (2019) and Marín-Lopez *et al*. (2019). Acute Alphaviruses infection and high mortality rate of immunocompromised mice were also demonstrated by Seymour *et al*. (2013) in O’nyong-nyong virus infection and detected in the current study. Regarding to virus tropism to different tissues, our data show higher virus titer on liver when compared to the brain and gonads, these results corroborate what was described in Marín-Lopez *et al*. (2019). In contrast to what has been demonstrated with MAYV in our study, the opposite is seen in the infection by ZIKV, witch replication is increased in organs such as brain and caused severe disease in the central nervous system (47,48).

The understanding of transmission process by mosquitoes and infection of the vertebrate host has relevance for public health, once arboviruses have been introduced in urban environments over the years and caused outbreaks. In the present study we described a model that can be used to evaluate vector competence of different arboviruses, pathogenesis in mammals infected similarly to natural viral transmission and also for future studies of viral genetic evolution. Grubaugh *et al*. (2017) used an birds animal to evaluate the replication, diversity and evolution of West Nile virus transmitted by *Culex* mosquitoes and showed a diversification and evolution of the virus in the different stages of the transmission cycle. Studies in this direction can help to elucidate the vector competence of some mosquito’s species and the adaptation of some viruses in specific environments (50).

Vector competence studies have been published for decades, however many works show insufficient data, lacking in detail or discrepant conclusions. In studies with vectors and vector-pathogen interaction it is important to focus on standardization of methods and nomenclature (29). The study design described here has the potential to be widely used with mosquitoes and arboviruses and reinforces the importance of conducting vector competence studies with complete transmission cycle. An animal model also expands the possibility of evaluating more generalist mosquito species in blood preference and are not easily artificially fed. In addition to studies of infection by one virus, our model can also be used in co-transmission and coinfection assays. A detailed understanding of how these scenarios impact virus transmission, as well as pathogenicity, is essential due to the current simultaneous co-circulation of several arboviruses in Brazil and other endemic countries.

## Conclusion

In conclusion, we demonstrate here a new model of MAYV transmission using mosquitoes and a vertebrate host and the species *Ae. aegypti* showed the ability to infect susceptible mice that developed disease, which were able, to infect another group of naive *Ae. aegypti* mosquitoes after seven days of infection. This animal model can be used with different arboviruses and mosquito species for a better understanding of the transmission cycle.

## Acknowledgments

We would like to thank Dr. Roberta Bronzoni from UFMT for kindly provided MAYV strain, MSc. Rafael Rosa for his help and training with the mice and Dr. George Diniz for his help with statistics analysis. We would also like to thank the team at the Entomology department for providing the mosquitoes.

